# An anti-inflammatory activation sequence governs macrophage transcriptional dynamics during tissue injury

**DOI:** 10.1101/2021.09.28.462132

**Authors:** Nicolas Denans, Nhung T. T. Tran, Madeleine E. Swall, Daniel C. Diaz, Jillian Blanck, Tatjana Piotrowski

## Abstract

Macrophages are essential for tissue repair and regeneration. Yet, the molecular programs, as well as the timing of their activation during and after tissue injury are poorly defined. Using a high spatio-temporal resolution single cell analysis of macrophages coupled with live imaging after sensory hair cell death in zebrafish, we find that the same population of macrophages transitions through a sequence of three major anti-inflammatory activation states. Macrophages first show a signature of glucocorticoid activation, then IL10 signaling and finally the induction of oxidative phosphorylation by IL4/Polyamine signaling. Importantly, loss-of-function of glucocorticoid and IL10 signaling shows that each step of the sequence is independently activated. Our results provide the first evidence that macrophages, in addition to a switch from M1 to M2, sequentially and independently transition though three anti-inflammatory pathways *in vivo* during tissue injury in a regenerating organ.

**One-Sentence Summary:** We show that macrophages are sequentially activated by three different anti-inflammatory pathways during tissue injury.

## Main Text

Innate immune cells, and more particularly macrophages, are essential during vertebrate embryonic development, tissue repair and regeneration (*1, 2*). Ablating macrophages during, or after injury of major organs in regenerating species blocks the regeneration process (*3-6*). Yet, the sequence of signals that controls their activity and their dynamics at a high spatio-temporal resolution are still poorly defined. Mammals can regenerate organs and tissues without scarring during embryonic and early postnatal stages but loose this ability as adults. A recent study in a mouse model of spinal cord injury showed that the crucial difference between neonatal mice and adults resides in differences in the activation state of macrophages (*7*). Strikingly, transplanted young macrophages allows adult mice to fully regenerate their spine. This stresses the urgent need to obtain a better understanding of embryonic macrophage activation states to develop therapeutics for adult regeneration.

Macrophages exist in various molecular activation states. They are commonly classified as pro-inflammatory (M1) or anti-inflammatory (M2) depending on the type of cytokines/signals they get activated with (*8*). However, this classification is an oversimplification. M1 macrophages can be activated by several pro-inflammatory signals such as Interleukin1, Interferon-gamma, lipopolysaccharide (LPS), while M2 are triggered by anti-inflammatory Glucocorticoids, Interleukin10, Interleukin4/13 and more (*9*). In addition, each of these signals triggers distinct downstream gene regulatory networks depending on the context and can have distinct functions. Nevertheless, a switch from M1 to M2 has been observed in different contexts of tissue injury (pathogen-induced-, sterile- or mixed injury). Two models have been proposed and observed in non-regenerative species: a “phenotypic switch”, where the same population switches from pro-to anti-inflammatory (*10-15*) or the “independent recruitment” of pro- and anti-inflammatory populations sequentially (*16-18*). Which model is at play during tissue regeneration is still poorly documented. A recent study in a model of tailfin regeneration in zebrafish demonstrated that the same macrophages can switch from pro-inflammatory (*tnfa*+) to anti-inflammatory (*cxcr4b*+) during the course of regeneration (*19*) favoring the phenotypic switch model. However, if cells can sequentially transition between several pro- or anti-inflammatory states during tissue injury is understudied mainly due to the lack of high spatio-temporal resolution analyses. Indeed, the majority of injury paradigms studied thus far occurs over many days, making it challenging to observe fast transitions between different macrophage activation states. Determining both the timing of the transitions as well as the genetic programs triggered by each activation state during tissue regeneration will be unvaluable to design targeted immunomodulatory therapies.

Here we took advantage of the rapid sensory hair cell (HC) regeneration that occurs in the zebrafish lateral line (*20, 21*) to uncover the molecular “What” (signals and genetic program) and “When” (sequence of activation) that control macrophage activity and dynamics during injury of a regenerating organ in zebrafish. This regenerating lateral line sensory system is powerful for several reasons: (i) antibiotic/neomycin treatment induces rapid HC death within minutes (*21*); (ii) HC regeneration occurs within hours and the first new pair of regenerated HCs is detected five hours after neomycin treatment (*22*); and (iii) the optical clarity of the larvae allows us to follow the behavioral dynamics of macrophages using confocal microscopy at high spatio-temporal resolution. Together, these characteristics enabled us to interrogate the transcriptional and behavioral dynamics of macrophage activation at an unprecedented spatio-temporal resolution.

## Results

### The same population of effector macrophages resolves inflammation within five hours in neuromasts

Tissue injury triggers an inflammatory response that involves the recruitment of macrophages to the injury site. These macrophages will be responsible for clearing cellular debris, remodeling the extracellular matrix and in some cases, provide signals to the tissue stem cells to start the repair process (*23*). We designated this population as ‘effector macrophages’ (*24*). It is not clear if effectors represent a single, or several macrophage populations. A recent study of muscle regeneration in zebrafish showed that two populations of macrophages were dynamically regulated after injury (*25*). One population reacted rapidly to tissue injury and left the damaged area within twenty-four hours, while a second population stayed in contact with the muscle stem cells for a longer period. Thus, the first step in understanding how effectors are molecularly regulated is to describe their dynamic after HC death. To follow both macrophages and HCs over time we study *Tg(mpeg:GFP*) and *Tg(she:lckmScarletI*) transgenic larvae that label macrophages and neuromast cells, respectively (Fig1A). Neuromast HCs are superficially located all along the larval body (Fig1A, movieS1). Treatment with neomycin for 30 min rapidly kills HCs by caspase-independent cell death (movieS2) (*26*). Macrophages start phagocytosing dead HCs as early as 15 min after the first cells die (movieS2-S3) (*27*). To describe the dynamics of effector macrophages from the time they enter the neuromast to when they leave during HC regeneration, we performed a macrophage recruitment assay. We treated the larvae with neomycin for 30 min and imaged both macrophages and neuromasts 1H, 3H, 5H and 7H after treatment (Fig1B, movieS4). Quantification of the number of macrophages surrounding and inside the neuromasts shows a rapid recruitment of effectors with a peak at 1H after neomycin (Fig1C-D). Subsequently, cells progressively return to homeostatic levels which they reach at 7H after neomycin treatment. Thus, there is a window of five hours when effector macrophages interact with neuromasts, which coincides with the appearance of the first regenerated HCs (*22*).

**Fig. 1.**
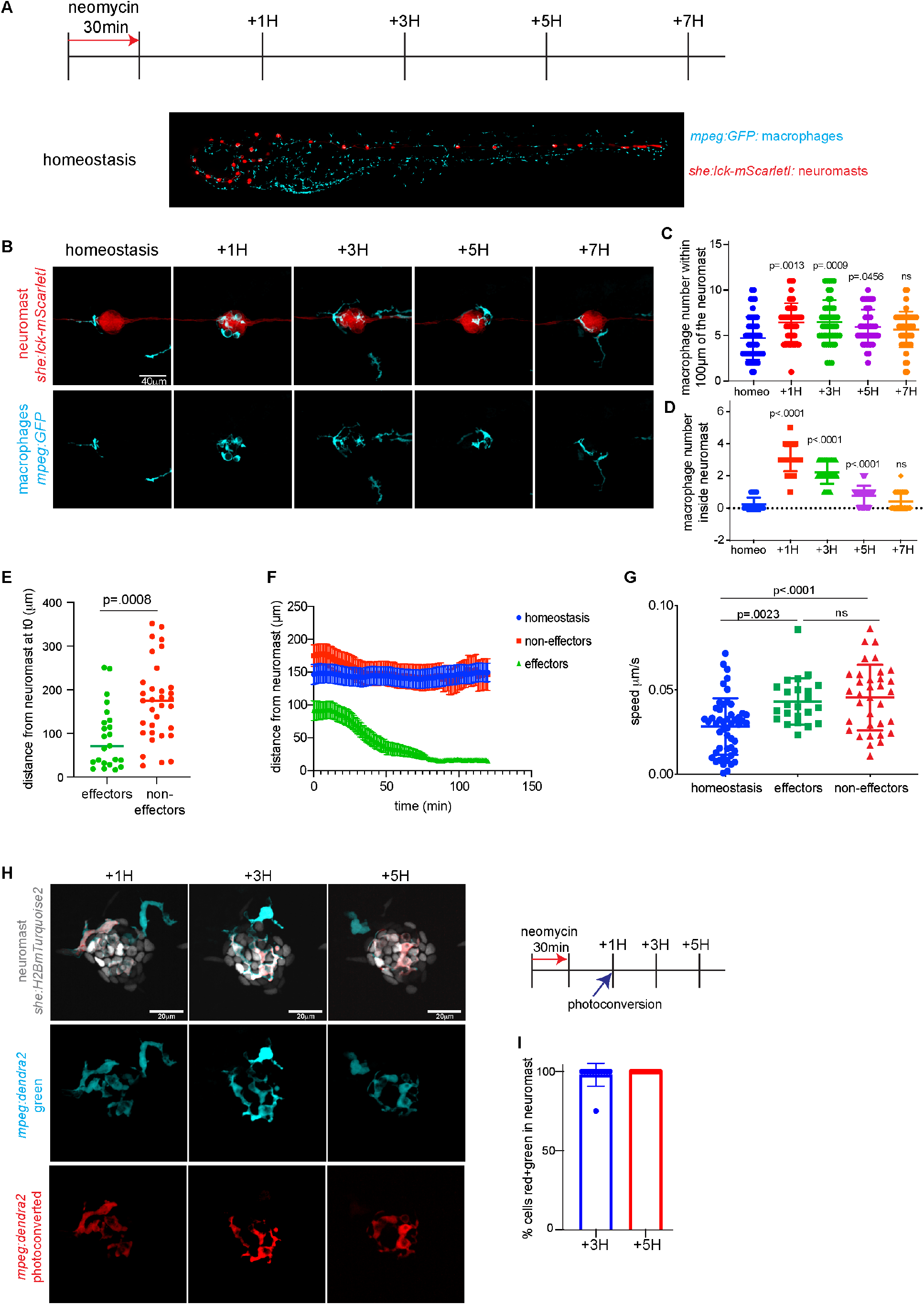
The same population of macrophages resolves HC death-induced inflammation within a five-hour window. (**A**) Schematics of neomycin regime and timepoint collections for the macrophage recruitment assay in the *Tg(she:lck-mScarletI;mpeg:GFP*) larvae. (**B**) Representative confocal images (projection of a 30mm z-stack) for the macrophage recruitment assay. (**C-D**) Quantification of macrophages in (C) or around (D) the neuromast. Each dot represents the number of macrophages per neuromast (3 neuromasts per larvae and 16 larvae per condition). P-values represent a post-hoc (Tuckey) test from each condition relative to homeostasis. (**E**) Quantification of the distance from the neuromast prior to HC death of effector and non-effector macrophages. Each dot represents a macrophage (8 larvae per condition). P-value represents a Student t-test. (**F**) Quantification of the distance from the neuromast over-time after HC death of effector and non-effector macrophages. Each dot represents a macrophage (8 larvae per condition). (**G**) Quantification of the macrophage velocity of effectors and non-effectors macrophages. Each dot represents a macrophage (8 larvae per condition). P-values represent a post-hoc test between each condition. (**H**) Representative confocal images (projection of a 30mm z-stack) for the macrophage photoconversion assay. (**I**) Quantification of the ratio of photoconverted effector macrophages over time. Each dot represents the percentage of macrophages per neuromast (1 neuromast per larvae and 12 larvae per condition).

To decipher if effector macrophages represent a specific population, we performed time-lapse recordings of macrophages and neuromasts during neomycin treatment and homeostasis in a large area of the larval trunk to identify where the effectors reside prior to HC death (movieS5). Quantification of the location of effector and non-effector macrophages before HC death shows that effectors are located closer to the neuromast than non-effectors (71μm+/-14μm vs 175μm+/-15μm, respectively) (Fig1E). In contrast to non-effectors, effectors show a rapid, directional migration toward the neuromasts in response to HC death (Fig1F). Interestingly, non-effector cells show an increase in non-directed cell velocity after neomycin treatment suggesting that HC death initially triggers a global injury response (Fig1G).

To assess if a single or multiple populations of effectors are recruited to neuromasts during the five hour window, we specifically photoconverted effector cells in neuromasts 1H after neomycin and quantified the ratio of photoconverted vs non-photoconverted macrophages 3H and 5H after HC death (Fig1H). Photoconverted cells represent an average of 98%+/-2% and 100% of the macrophages inside neuromasts at 3H and 5H after HC death, respectively, demonstrating that a single population of effectors is recruited during the five hour window (Fig1I).

### High temporal resolution scRNA-seq identifies a population of effector macrophages

To molecularly characterize all macrophage populations, identify the effector population and characterize their activation sequence and underlying molecular program over the course of HC death, we performed a scRNA-seq time course of *Tg(mpeg:GFP*) transgenic larvae. We dissociated 600 5dpf larvae each during homeostasis, and 1H, 3H and 5H after a 30min neomycin treatment and FACsorted GFP+ cells for 10x Chromium genomics scRNA-seq (Fig2A). To determine the earliest activation of macrophages we also collected GFP+ cells immediately after a 15 min treatment with neomycin (Fig2A). We integrated these five timepoints using Seurat and performed UMAP dimensional reduction (Fig2B). We downsampled the number of cells per timepoint to fourteen thousand to avoid a bias based on over-representation of a specific timepoint (FigS1A). The seventy thousand cell atlas of all *mpeg:*GFP cells of 5 dpf larvae revealed that *mpeg*:GFP, in addition to macrophages, also labels several other immune cell types, some of which have also been recently observed in 4dpf larvae (*28*). We identified dendritic-like cells (DC-like) based on their expression of the master regulator *flt3 (29*), as well as *spock3* and *hepacam2* (Fig2B, FigS1B-C and TableS1). Previously described antigen presenting metaphocytes (*30*) are marked by *cldnh, epcam* and *prox1a* (Fig2B, FigS1B-C and TableS1). Two clusters of natural-killer like cells express their master regulator *eomesa* (*31*) (NK-like) and *gata3* (*32*) (NK-like2) (Fig2B, FigS1B-C and TableS1). A neutrophil population is labeled by *mpx* and *lyz (33*), an unidentified population expresses skin (*rbp4, sparc*) and collagen markers (*col1a2*) as well as two unidentified small clusters labelled by *kng1* and *acta3b* respectively (Fig2B, FigS1B-C and TableS1). Our analysis revealed an unexpected heterogeneity of the macrophage population consisting of eight clusters that cluster closely together (Fig2B). Proliferating macrophages form a cluster characterized by *pcna, mki67* and *tubb2b* (Fig2B, FigS1B-C and TableS1). Another cluster that is heavily influenced by ribosome genes we called the ‘translation’ cluster (Fig2B, FigS1B-C and TableS1). Cells in the small ‘stat1b’ cluster show an Interferon signaling response signature (*stat1b, isg15, cxcl20*) (Fig2B, FigS1B-C and TableS1) that is also induced in response to an endemic picornavirus in zebrafish facilities (*34*). Another cluster is marked by genes classically upregulated in response to bacteria (*35, 36*) that we named ‘irg1/acod1’ (*irg1/acod1, hamp, mxc*) (Fig2B, FigS1B-C and TableS1). In addition, we identified a cluster that did not show unique markers but is broadly labelled by *f13a1b, junba* and *btg1* (called ‘fa13a1b’) (Fig2B, FigS1B-C and TableS1); a cluster that expresses markers of potentially immature microglia, such as *mcamb, apoeb* (*37*) and *apoc1* (called ‘mcamb’) (Fig2B, FigS1B-C and TableS1); and two clusters that represent unidentified macrophage states or populations expressing *runx3, cxcl19, cxcl8a* (called ‘runx3’) and *tspan10, slc43a3b, illr1* (called ‘tspan10’), respectively (Fig2B, FigS1B-C and TableS1).

To characterize macrophage clusters that show transcriptional changes during the neomycin time course, we performed differential gene expression for each cluster at each timepoint. Quantification of the numbers of up-and down-regulated genes during the time course shows that several macrophage clusters respond to neomycin treatment (Fig2C, D, H, L and FigS2A-B). To determine which macrophages enter the neuromasts and represent effector cells, we performed fluorescent *in situ* hybridization using the hybridization chain reaction (HCR-FISH (*38*)) with the macrophage cluster markers *irg1*/*acod1, f13a1b, mcamb, tspan10, runx3*, as well as the non-macrophage NK-like (*eomesa*) and DC-like (*hepacam2*) clusters as negative controls (Fig2D-O, FigS2A-D’’). HCR-FISH was performed on *mpeg:GFP* larvae 1H after neomycin, when all effector cells have migrated into the neuromasts. This screen demonstrated that 48% of the effectors are labelled with *irg1*/*acod1* (Fig2D-G), 18% with *f13a1b* (Fig2H-K) and 34% with *mcamb* (fig2L-O). The *tspan10* marker did not label any effector cells (FigS2A-A”), while we found that 3% of the cells were labeled with *runx3* (FigS2 B-B”). As expected, no effector cells were labelled with *eomesa* (NK-like cells) (FigS2 C-C”) or *hepacam2* (DC-like cells) (FigS2 D-D”). Additionally, we performed a macrophage recruitment assay with two newly generated transgenic reporter lines that drive the expression of a red fluorescent protein under the promoter of *irg1*/*acod1* and *stat1b*. These experiments confirmed that ‘irg1/acod1’ cells are indeed effector cells, whereas ‘stat1b’ cells do not enter neuromasts (FigS3A-D). Altogether we conclude that the effector population is composed of cells belonging to the ‘irg1/acod1’, ‘f13a1b’ and ‘mcamb’ clusters and focused our analysis on these cells as effector macrophages.

**Fig. 2.**
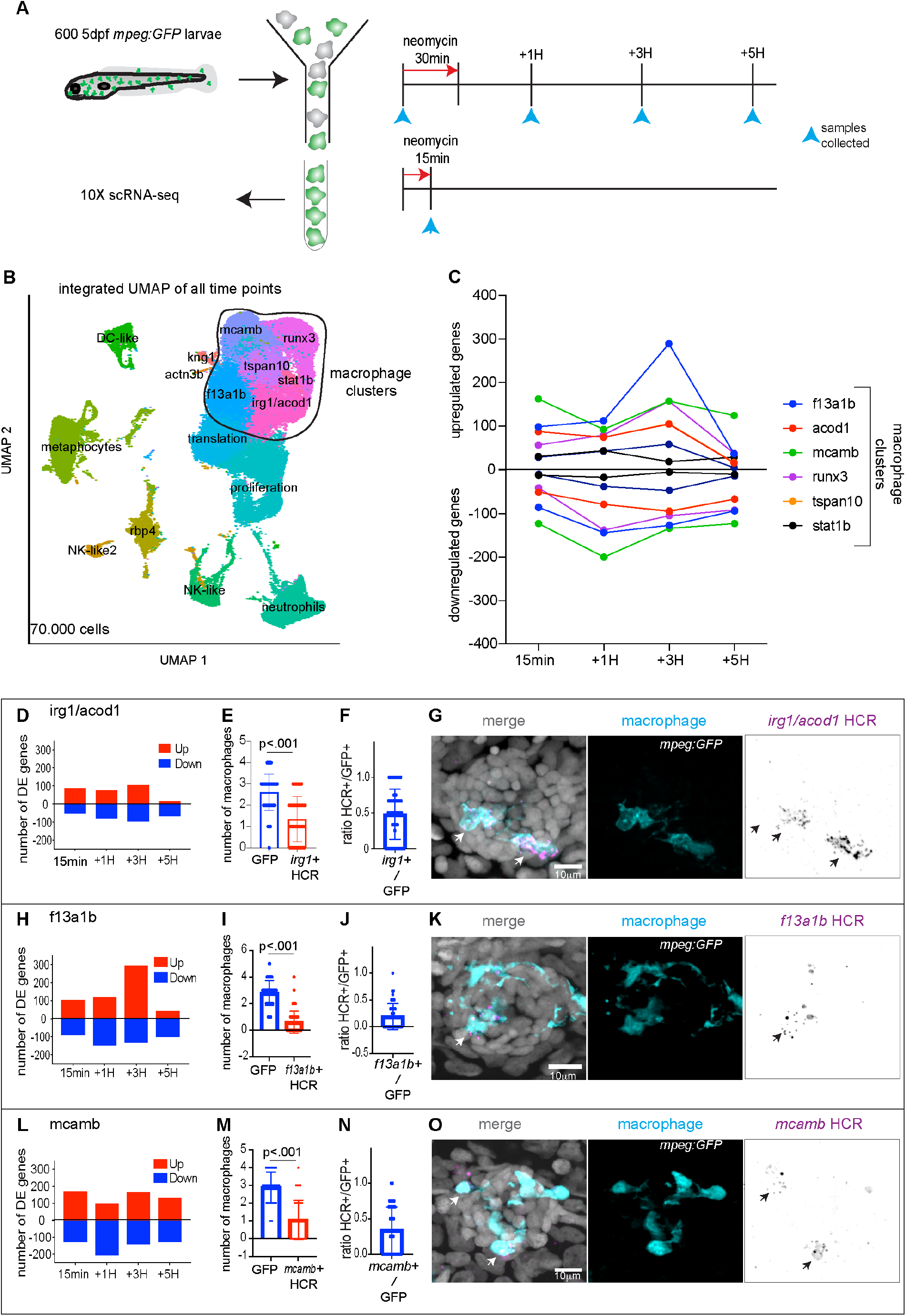
Combination of scRNA-seq and HCR-FISH identifies a population of effector macrophages during HC regeneration. **(A**) Schematics of neomycin regime and timepoints collection for scRNA-seq. (**B**) Integrated UMAP of the five timepoints. Cluster names are labelled on the UMAP. Macrophage clusters are circled. (**C**) Quantification of genes differentially up- and downregulated at each timepoint within the macrophages clusters. (**D, H, L**) Quantification of genes differentially up- and downregulated at each timepoint for (D) the ‘irg1/acod1’, (H) ‘f13a1b’ and (L) ‘mcamb’ clusters. (**E, I, M**) Quantifications of GFP+ effector cells and HCR effector cells for (E) *irg1/acod1*, (I) f*13a1b* and (M) *mcamb*. Each dot represents the number of macrophages per neuromast (5 neuromasts per larvae and 16 larvae). P-values represent non-parametric Student t-test. (**F, J, N**) Quantifications of the ratio between HCR+ cells and GFP+ effector macrophages in the (F) *irg1/acod1*, (J) f*13a1b* and (N) *mcamb* clusters. (**G, K, O**) Representative confocal images (projection of a 30μm z-stack) of HCR-FISH within the effector macrophages for (G) *irg1/acod1*, (K) f*13a1b* and (O) *mcamb*.

### An anti-inflammatory activation sequence in macrophages during HC death

To identify the core molecular programs driving the effector macrophages activation states at each timepoint, we compared the differentially expressed genes of the three clusters and analyzed the genes that were shared (TableS2). We detected that at the fifteen minutes neomycin timepoint the Glucocorticoid (GR) pathway targets (*arl5c, jdp2b, dusp1, klf9, tsc22d3, nfkbiab, sgk1, fkbp5*) (*39, 40*) are strongly and transiently upregulated (Fig3A, S4A, TableS2). HCR-FISH for *dusp1* revealed that this activation of the GR pathway is systemic (Fig3B). Likewise, *interleukin10 receptor alpha* (*il10ra*) is also strongly upregulated (Fig3A, S4A, TableS2). We confirmed the expression of *il10ra* in 75% of effector cells by HCR-FISH (Fig3C-D). Gene ontology and pathway analyses using Metascape (*41*) showed an enrichment for genes involved in regulation of apoptosis (*gadd45ab, gadd45bb, xiap, gbp, btg2, ddit3 and mcl1a*) and GR pathway activation (TableS2) within the upregulated genes, while ‘oxidative phosphorylation’ is the most enriched term within the downregulated genes at the 15 min timepoint (FigS4B, TableS2). Other markers of anti-inflammatory macrophages that have not been linked to GR activation, such as *cxcr4b, irf2, lpn1, mknk2* and *ets2* (*42-46*) are also strongly upregulated (FigS4A, TableS2). Of note, *ncf1* and *nrros* (*47, 48*), both negative regulators of reactive oxygen species, are strongly upregulated (FigS4A, TableS2). Unexpectedly, our stringent analysis did not reveal a robust upregulation of pro-inflammatory cytokines and the pro-inflammatory *il1b* is only upregulated in the ‘irg1/acod1’ cluster at the 15 min neomycin timepoint, while no transcriptional upregulation of *tnfa, il6* or *il12a* could be detected (Fig3A and S5). This finding suggests that a transition from a pro-inflammatory (*il1b*+) to an anti-inflammatory (GR+) state occurs rapidly within minutes after HC death. The immediate upregulation of the *il10ra* receptor at 15 min leads to the activation of IL10 target genes activation at 1H after neomycin (*socs3b, ccr12b*.*2, pim1* and *fgl2a*) (*49-51*) (Fig3A), which follows the increase in *il10ra* receptor expression at the prior timepoint. HCR-FISH confirms that 80% of the effector cells express *fgl2a* (Fig3E-F). Other master activators of anti-inflammatory macrophage states, such as the *interleukin4 receptor il4r*.*1* and the rate limiting enzymes of the polyamine pathway, *odc1* and *smox*, are also upregulated (Fig3A, S4A, TableS2). At 3H, in addition to the inhibition of pro-inflammatory cytokines by IL4, the combination of Polyamine and IL4 signaling induces oxidative phosphorylation in macrophages (*52, 53*) (Fig3A, S4A, tableS2). For example, genes belonging to the mitochondria respiratory chain complexI (*ndufab1b, ndufb3, mt-nd1*), complexIII (*cycsb, uqcr10, uqcrfs1, uqcrh*), complexIV (*cox17, cox7a2a, cox7c, cox7b*) and complexV (*atp5g1, atp5g3a, atp5j, mt-atp6*) are induced (Fig3A, S4A, TableS2). Likewise, Pathway and Gene Ontology (GO) analysis shows ‘Oxidative phosphorylation’ as the most enriched term (TableS2). Interestingly, *manf*, a gene required for retina repair and regeneration in fly and mouse (*54*) is strongly upregulated at the three-hour timepoint hinting toward a switch to a repair state of macrophages (Fig3A, S4A). In contrast to the earlier timepoints, very few genes are found specifically upregulated at the 5H timepoint (Fig3A, S4A). *grn2*, a pro-granulin growth factor regulator of the anti-inflammatory macrophage phenotype (*55*) is highly upregulated. Interestingly, in *grn* knock-out mice, muscle injury leads to a persistence of macrophages at the injury site suggesting a role for *grn* in regulating macrophage dynamics (*56*). Altogether, this analysis reveals a linear sequence of macrophage anti-inflammatory activation immediately after HC death starting with Glucocorticoid signaling, followed by IL10 signaling, and lastly a combination of Polyamine and IL4 signaling, which induces oxidative phosphorylation at the transcriptional level. To our knowledge, this is the first report of an *in vivo* linear sequence of three major anti-inflammatory activation pathways after injury. This suggests that in addition to a transition from a pro-inflammatory M1 to an anti-inflammatory M2 state, the same macrophages transition through different anti-inflammatory states to potentially regulate their dynamics/activity.

**Figure 3.**
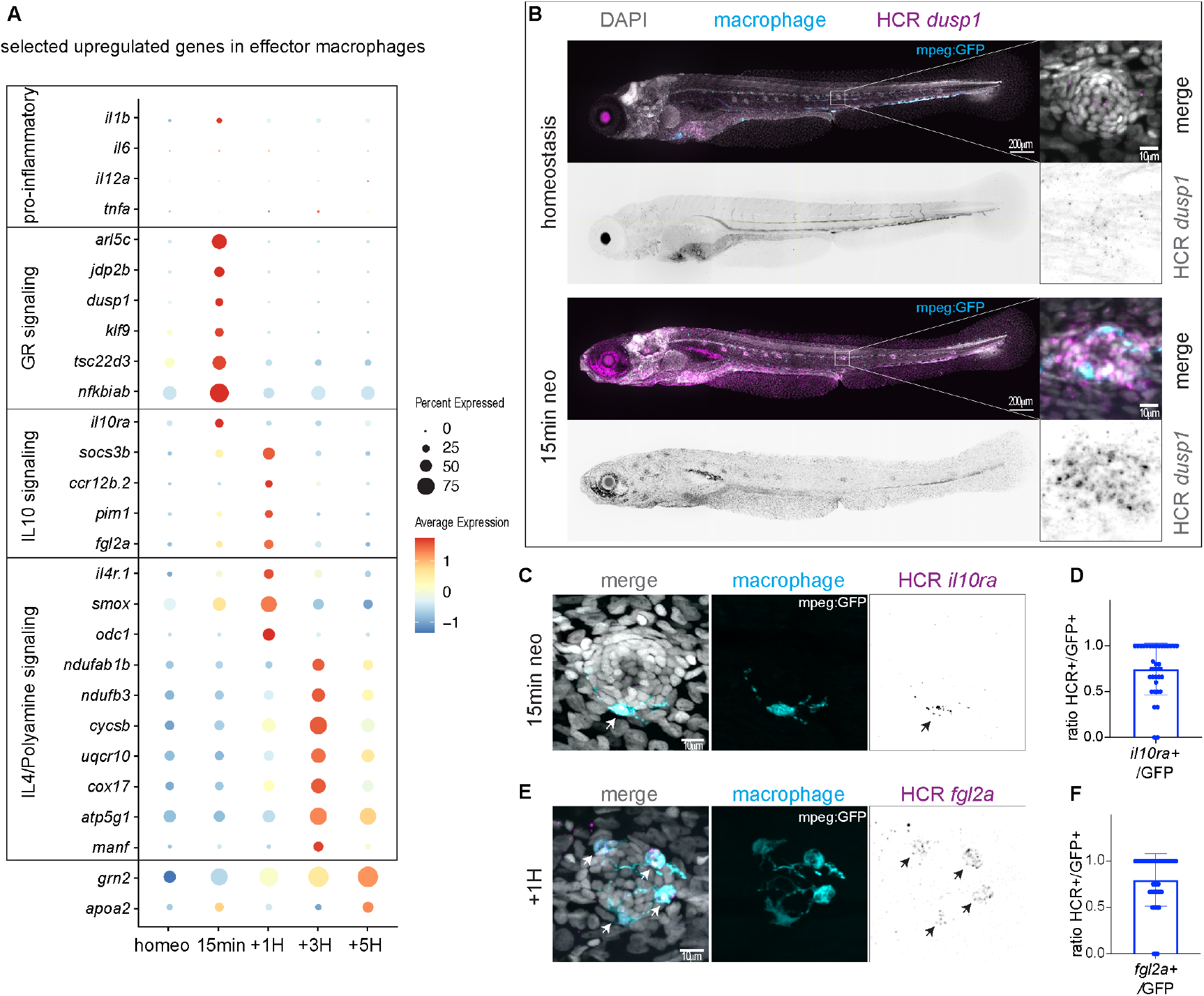
Transcriptional dynamics reveal an anti-inflammatory activation sequence in effector macrophages. **(A)** Dot-Plot of selected differentially upregulated genes for each timepoint of the scRNA-seq time course within the effector population. (**B**) Representative confocal images (Maximum projection of a 200μm z-stack) of an HCR-FISH for *dusp1* in a 5dpf larvae (left) and zoom image of a representative neuromast (right). (**C, E**) Representative confocal images (projection of a 30μm z-stack) of HCR-FISH within the effector macrophages for (C) *il10ra*, (E) f*gl2a*. (**D, F**) Quantifications of the ratio between HCR+ cells and GFP+ effector macrophages for (D) *il10ra*, (F) f*gl2a*. (12 larvae with 3 neuromasts per larva).

### Each anti-inflammatory state is independently activated

The discovery of this linear anti-inflammatory sequence of effector macrophage activation raises the interesting question if epistatic relationships between Glucocorticoid signaling and IL10 signaling and between IL10 signaling and IL4/Polyamine signaling exist.

To address this question, we first inhibited Glucocorticoid signaling during HC death using the GR inhibitor RU486. HRC-FISH for the GR target *dusp1* confirmed that a 10μM treatment with RU486 was efficient in blocking GR activation (Fig4A). HCR-FISH for *il10ra* at the 15 min timepoint, as well as for the IL10 signaling target gene *fgl2a* at the 1H timepoint showed no downregulation of these genes in effector macrophages after RU486 treatment (Fig4B-E). This demonstrates that GR activation is not required for IL10 activation in effector macrophages.

**Figure 4.**
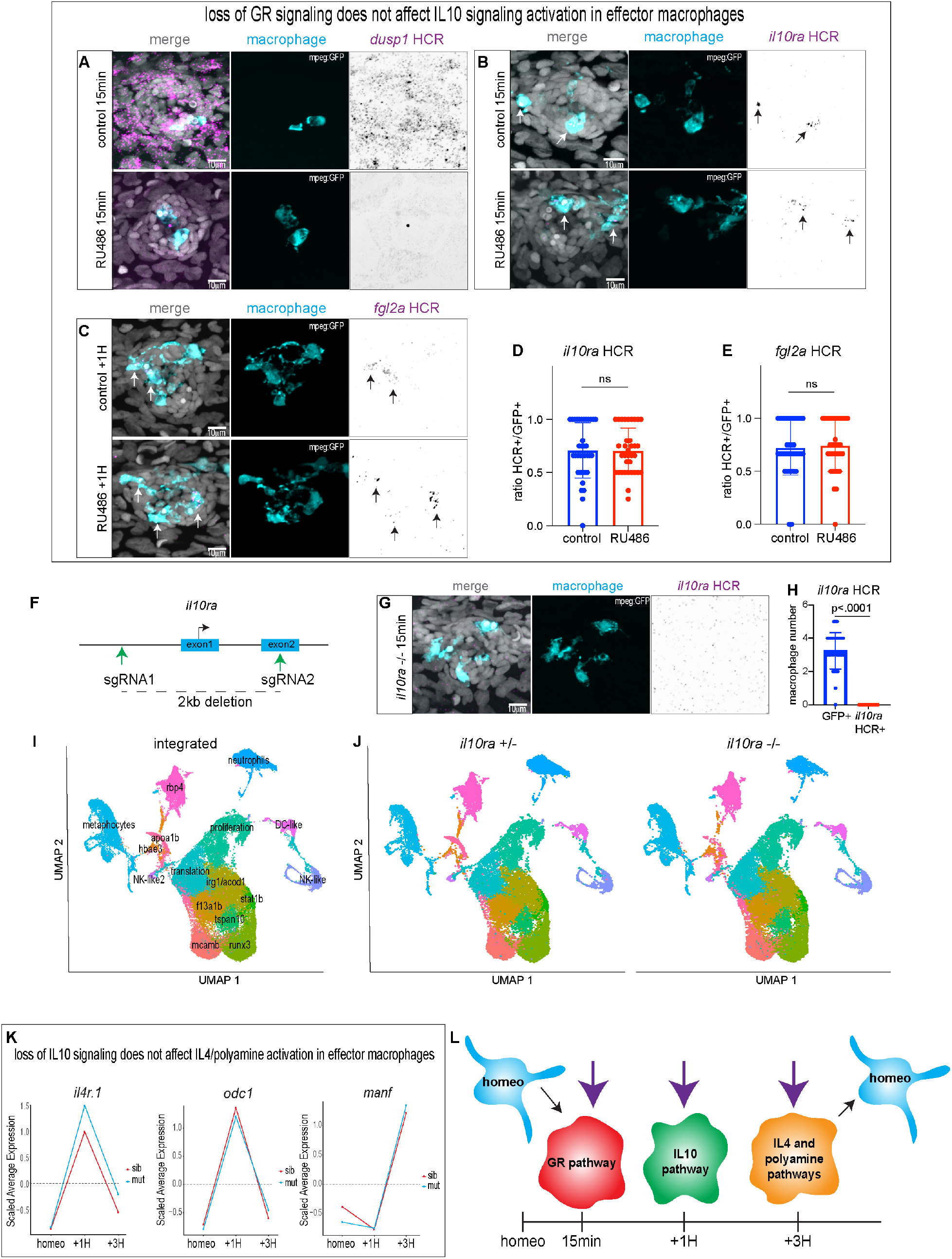
Each anti-inflammatory state is independently activated. **(A)** Representative confocal images (maximum projection of a 30μm z-stack) of an HCR-FISH for *dusp1* in a neuromast (10 larvae per condition and 30 neuromasts total). (**B, C**) Representative confocal images (projection of a 30μm z-stack) of HCR-FISH within the effector macrophages for (B) *il10ra* and (C) f*gl2a* (12 larvae per conditions and 36 neuromasts total) (**D, E**) Quantifications of the ratio between HCR+ cells and GFP+ effector macrophages for (D) *il10ra* and (E) f*gl2a*. (**F**) Schematics representing the CRISPR/Cas9 design used to generate the *il10ra* mutant zebrafish.(**G**) Representative confocal images (projection of a 30μm z-stack) of HCR-FISH within the effector macrophages for *il10ra* in the *il10ra* homozygous mutant. (**H**) Quantifications of GFP+ effector cells and HCR in effector cells for *il10ra* in the *il10ra* homozygous mutant. Each dot represents the number of macrophages per neuromast (33 neuromasts from 11 larvae). P values represent non-parametric Student t-test. (**I**) Integrated UMAP of the six datasets for the *il10ra* mutant. Cluster names are labelled on the UMAP. (**J**) Split UMAP per condition (*il10ra*+/- and *il10ra*-/-). (**K**) Line plots representing the average expression for each timepoint between the *il10ra* mutant (cyan) and the sibling (red) from the scRNA-seq integrated dataset. (**L**) Model of independent activation of the three anti-inflammatory pathways in effector macrophages during the HC regeneration time course. Resting macrophages are represented in cyan. Purple arrows represent the independent induction of each macrophage activation state.

To test for a possible epistatic relationship between IL10 signaling and the following activation state (Polyamine + IL4 signaling), we generated a mutant for *il10ra* by deleting the promoter region, as well as the first exon using CRISPR/Cas9 (Fig4F). HCR-FISH for *il10ra* in homozygous mutants confirmed the absence of transcript (Fig4G-H). We also performed HCR-FISH for the target gene *fgl2a* at the 1H timepoint and found a strong downregulation in effector cells in the mutant (64%+/-7% in homozygous vs 21%+/-5% in heterozygous) (FigS6A-B). Next, we performed a scRNA-seq time course after neomycin treatment in homozygous and heterozygous *il10ra* larvae (FigS7A). Integration of these six datasets (sixty thousand cells) using Seurat and UMAP dimensional reduction shows that the effector clusters (‘irg1/acod1’, ‘mcamb’ and ‘f13a1b’) are conserved in mutant larvae (Fig4I-J, S7B-C, TableS3). Likewise, quantification of the expression levels and dynamics of *il4r*.*1*, and *odc1* shows that they are unaffected in the mutants compared to the siblings (Fig4K). The subsequent activation of genes related to oxidative phosphorylation and *manf* 3H after neomycin is also unaffected (Fig4K and S8). Thus, the induction of oxidative phosphorylation by IL4/Polyamine signaling is independent of IL10 signaling activity. This important result reveals that, while a sequential induction of anti-inflammatory pathways underlies macrophage activation during HCs regeneration, its components are independently activated (Fig4L). Therefore, activation of a single anti-inflammatory pathway is likely not sufficient to induce proper tissue regeneration and a sequential and independent activation of the three pathways might be required.

## Discussion

Macrophage molecular activation regulates the dynamic and activity of these phagocytes and modulation of their activation states and their dynamic can have dramatic effects on tissue repair and regeneration (*57*). Here, using a high spatio-temporal resolution analysis of macrophage activation during HC regeneration, we provide evidence that the same population of macrophages is sequentially and independently activated by three major anti-inflammatory pathways.

It is now broadly documented that macrophages adopt a pro-inflammatory activation state immediately after injury (*19, 28, 58-60*). The role for this pro-inflammatory phase is mainly to attract more macrophages to the injury site if the organ does not possess a resident population, or if the size of the injury requires more macrophages (*61*). This first phase must be followed by an anti-inflammatory phase to resolve the inflammation and ensure proper tissue repair. Our analysis shows that in the lateral line, a single population of tissue resident macrophages resolves HC death induced inflammation, favoring the phenotypic switch model. Furthermore, our data show a very short pro-inflammatory phase and a rapid transition to an anti-inflammatory state marked by the strong and systemic activation of the GR pathway. A possible explanation for this lack of a strong pro-inflammatory activation of macrophages after HC death is that the tissue resident population is sufficient to resolve inflammation and the recruitment of inflammatory macrophages is not required. This hypothesis is supported by our finding that the effector macrophages reside in immediate vicinity of the neuromasts during homeostasis and that after HC loss on average only three macrophages are detected in neuromasts.

The anti-inflammatory role of the GR pathway has been extensively documented (*39, 62-64*). It can both directly inhibit pro-inflammatory gene transcription by direct binding of the GR receptor to their enhancers/promoters or by tethering the pro-inflammatory activators AP-1 and Nfkb (*64*). GR also triggers a rapid and large-scale chromatin unwinding, potentially creating a permissive environment for drastic changes in gene expression required for macrophage activity (*65*). Therefore, the strong and transient activation of the GR pathway immediately as the first HCs start to die likely turns off the transcription of pro-inflammatory cytokines.

The short phase of GR activation is immediately followed by IL10 signaling activation and a subsequent transition to a IL4/Polyamine activation state that induces oxidative phosphorylation. A recent study in mouse Bone Marrow Derived Macrophages (BMDM) showed that part of the IL10 anti-inflammatory function is to inhibit glycolysis while promoting oxidative phosphorylation after treatment with LPS, which mimics a bacterial infection (*66*). Our *il10ra* mutant analysis demonstrates that lack of IL10 signaling after HC death does not affect the induction of oxidative phosphorylation. Co-stimulation of mouse BMDM with LPS and IL10 also leads to the upregulation of the IL4 receptor (*67*) whereas the loss of IL10 in zebrafish does not affect the induction of *il4r*.*1*. These discrepancies suggest that IL10 and IL4 signaling are differently regulated in response to bacterial infection versus tissue injury.

The IL4/OXPHOS state is characteristic for wound healing macrophages and responsible for the last step of inflammation resolution (*2, 68-70*). In addition, the induction of the pro-repair gene *manf*, which is required for retina regeneration (*54*), suggests that at the 3H timepoint macrophages transition to a wound-healing state. Altogether, our *in vivo* data demonstrate the sequential and independent transition of effector macrophages through three major anti-inflammatory states. This finding has important implications for the design of targeted immunomodulatory therapies.

## Acknowledgments

We thank the Piotrowski lab members for insightful discussions and Dr. Mark Lush, Julia Peloggia, and Daniela Münch for critical reading of the manuscript. We thank Allison Scott for technical assistance. We are also grateful to the Stowers Institute Core Facilities (Aquatics team, Cytometry Core, Imaging Core, Molecular Biology Core) for their technical expertise.

## Author contributions

N.D. designed and performed the experiments, analyzed, and interpreted the data, and wrote the manuscript; N.T. analyzed and interpreted the data; D.D. analyzed and interpreted the data; M.S. performed experiments; J.B. performed experiments; T.P. designed the experiments, analyzed, and interpreted the data, and wrote the manuscript.

## Data availability

Original data underlying this manuscript can be accessed from the Stowers Original Data Repository at http://www.stowers.org/research/publications/libpb-1663

## Supplementary Materials

Materials and Methods

Figs. S1 to S8

Tables S1 to S3

Movies S1 to S5

