## Supplementary materials for "An anti-inflammatory activation sequence governs macrophage transcriptional dynamics during tissue injury"

**This PDF file includes:**

Materials and Methods
Figs. S1 to S8
Captions for Movies S1 to S2

**Other Supplementary Materials for this manuscript include the following:**

Movies S1 to S5
Tables S1 to S3

### Materials and Methods

Zebrafish lines and husbandry

Animal work followed the guidelines of the animal ethics committee (IACUC review board) at the Stowers Institute for Medical Research. The following zebrafish transgenic and mutant lines were used: *Tg*(*mpeg1.1:*EGFP)$^{gl22}$, abbreviated as *mpeg*:GFP (*1*); *Tg*(*she*:lckmScarletI)$^{psi70Tg}$; *Tg*(*she*:H2BmTurquoise2)$^{psi72Tg}$; *Tg*(*Myo6b*:lck-mScarlet-I)$^{psi67Tg}$ (*2*); *Tg*(-5.6*irg1*:lck-mScarletI/*acry*:mScarletI) $^{psi73Tg}$ referred as *irg1*:lck-mScarletI. 5.6kb directly upstream of *irg1* start codon were cloned from genomic DNA and drive the expression of lck-mScarletI.; *Tg*(-9*stat1b*:lck-mScarletI/*acry*:mScarletI) $^{psi75Tg}$ referred as *stat1b*:lck-mScarletI. 9kb directly upstream of *stat1b* start codon were cloned from genomic DNA and drive the expression of lck-mScarletI.; *Tg*(*she*:gap43-GFP) ) $^{psi74Tg}$; *Tg*(*mpeg1.1*:dendra2)$^{uwm12}$ referred as *mpeg*:dendra2 (kind gift from Dr. Anna Hunttenlocher (*3*)); All transgenic lines were generated using the tol2 system (*4*).

The *il10ra* $^{psi71}$ mutant was generated using CRISPR/Cas9. sgRNAs were selected using CRISPRscan (*5*) to delete a 2kb region from the putative promoter to the second exon. sgRNA1: 5-GTGTTTTGGTCGGGTGTGGT-3; sgRNA2: 5-GGTACGGGGGCTTCATTGGG-3. Genotyping primers to assess for the deletion are: Fwd: 5-TGCATAACAGCTCAGCCATCTTCTC-3; Rv: 5- CATGTCACATTCTTCCACAATGTTCC-3

Sensory Hair Cells Ablation

To ablate hair cells, 5dpf embryos were treated with 300µM neomycin (Sigma-Aldrich, St Louis, MO, USA) for 30min at 28 °C or for a 15minutes pulse. Following, embryos were washed with 0.5x E2 medium (7.5 mM NaCl, 0.25 mM KCl, 0.5 mM $MgSO_4$, 75 mM $KH_2PO_4$, 25 mM $Na_2HPO_4$, 0.5 mM $CaCl_2$, 0.5 mg/L $NaHCO_3$, pH = 7.4) and incubated at 28 °C until further experimental needed for further experiments.

Embryo dissociation and FACS

600 GFP-positive 5dpf larvae were anesthetized with tricaine for ~1 min until they stopped moving (1:20 dilution of 4g/L tricaine in 0.5x E2 medium). To dissociate the larvae, we placed them into 2 wells (300 larvae each) containing strainers (BD Falcon Cell Strainer (BD Biosciences, San Jose, USA), quickly rinsed the larvae in ice-cold DPBS and added 4.5 ml cold 0.25% trypsin-EDTA (Thermo Fisher Scientific, Waltham, USA) supplemented with 1uM ActinomycinD (SIGMA A1410). The larvae (300 each) were then transferred to one 5ml polypropylene conical tube (placed on ice) with a disposable transfer pipet. The larvae were dissociated by trituration with a Pasteur pipette hooked to a pipetboy until all GFP+ cells were dissociated (~7minutes) on ice. Cells in suspensions were first separated from the larval bodies by filtering the suspension through a Filcons 70 µm cell strainer (BD Biosciences, San Jose, CA, USA) into a 5ml polypropylene round-bottom tube. Subsequently, the cells were centrifuged at 2000 rpm (720 x g) for 5 min at 4°C. To wash off the trypsin-EDTA, we removed it, added ice-

cold DPBS with 1uM ActinomycinD and centrifuged the cells at 2000 rpm (720 x g) for 5 min at 4°C. Resuspended cells in fresh ice-cold DPBS with 1uM ActinomycinD were filtered through a Filcons 70 µm cell strainer (BD Biosciences, San Jose, CA, USA) into a falcon round bottom 5ml tube (snap cap tube, Falcon-Corning, Glendale, AZ, USA) and immediately FACSorted. Samples were sorted on a BD Influx with a 100µm tip at 20psi with 1X PBS, directly into chilled 90% MetOH. Prior to sorting, samples were stained with 25uM DRAQ5 for 5 minutes on ice. DRAQ5 is excited by a 647nm laser at 100mW with detection at 720/40nm. GFP is excited by a 488nm laser at 100mW with detection at 528/28nm. Single color controls were analyzed to confirm spectral compensation is unnecessary; the GFP+ gate was set using a non-expressing sample stained with DRAQ5 as an FMO (fluorescence minus one) control.

10X Chromium scRNA-seq library construction

Methanol fixed cells were rehydrated with rehydration buffer (1% BSA and 0.5 U/µl RNase-inhibitor in ice-cold DPBS). Approximately 20.000 cells were loaded into the Chromium Single Cell Controller (10x Genomics). For library preparation, Chromium Next GEM Single Cell 3' GEM, Library Gel Bead Kit v3.1 was used. The sample concentration was measured on a Bioanalyzer (Agilent) and sequenced with Novaseq6000 Kit v3 with read length of 28 bp Read 1, 8 bp i7 index and 91 bp Read 2 (150 cycles) (Illumina).

scRNA-Seq read alignment and quantification

Raw reads were demultiplexed and aligned to version 10 of the zebrafish reference transcriptome (danRer10, Ensembl release 91) following the 10X Genomics' CellRanger (v2.1.1) pipeline for the macrophage time course dataset. Prior to the downstream QC filtering we obtained the following cell numbers: 20,676 (homeostasis), 17,187 (15min), 29,846 (1h), 20,540 (3h), and 26,413 (5h) using CellRanger's cell-association algorithm. To normalize the cell numbers across the time course, we randomly subsampled 14,000 cells from each timepoint. Post-filtering, the number of cells per sample were 13,937 (homeostasis), 13,908 (15min), 13,921 (1h), 13,886 (3h), and 13,921 (5h). The mean number of genes per cell per sample were 1,475 (homeostasis), 1,306 (15min), 1,174 (1h), 1,558 (3h) and 1,488 (5h).

For the *il10ra* heterozygous (also referred to as sibling) versus homozygous (also referred to as mutant) scRNA-seq dataset, raw reads were demultiplexed and aligned to version 11 of the zebrafish reference transcriptome (danRer11, Ensembl release 102) following 10X Genomics' CellRanger (v6.0.1). Prior to QC filtering, we obtained the following cell numbers for the heterozygous (sibling) samples: 14,159 (homeostasis), 12,720 (1h), and 13,793 (3h) using CellRanger's cell-association algorithm. We obtained the following cell numbers for the homozygous (mutant) samples: 12,004 (homeostasis), 14,183 (1h), and 19,546 (3h) using CellRanger's cell-association algorithm. Post-filtering, the number of cells per sibling sample were 7,201 (homeostasis), 9,517 (1h), 10,705 (3h) and 7,895 (homeostasis), 10,806 (1hr), and 13,693 (3hr) for the mutant samples. The mean number of genes per cell per sibling samples were 2,073 (homeostasis), 1,347 (1h), and 1,560 (3h). The mean number of genes per cells per mutant samples were 1,825 (homeostasis), 1,248 (1h), and 1,628 (3h). All raw data for the macrophage time course and *il10ra* sibling versus mutant samples including sorted BAM files and count matrices produced by CellRanger has been deposited in Gene Expression Omnibus (GEO) database, www.ncbi.nlm.nih.gov/geo (accession no. GEOx).

Pre-processing, quality filtering and batch integration

To distinguish zebrafish repeated gene symbols with unique Ensembl IDs, we modified the count matrices outputted from the CellRanger pipeline using a custom R script (https://github.com/ntran95/SeuratExtensions/blob/main/R/makeuniq_updated.R). Each repeated gene symbol is annotated with an asterisk followed by an incrementing number. A comprehensive gene list with unique repeated symbol annotations can be downloaded from our interactive web applications (https://piotrowskilab.shinyapps.io/macrophage_timeseries_scRNAseq_pub_2021/, https://piotrowskilab.shinyapps.io/il10ra_sib_vs_mut_mphg_nd0897/). Since the macrophage time course and *il10ra* samples were aligned to different reference transcriptomes (Ensembl release 91 versus Ensembl release 102 respectively), we provide both gene symbol conversions in the supplemental gene lists to account for any differences between the two versions (TableS1, TableS3).

For the macrophage time course dataset, low quality cells or cells containing doublets with reads greater than 20,000, reads less than 600 and mitochondrial contamination greater than 5% for each sample were filtered from the subsequent analysis. Genes present in less than 20 cells were also removed from the dataset.

For the *il10ra* samples, low quality cells or cells containing doublets with reads greater than 50,000, genes per cell less than 500, genes per cell greater than 7,500 and mitochondrial contamination greater than 5% from each sample were filtered from the subsequent analysis. Genes present in less than 20 cells were also removed from the dataset.

Both datasets were integrated following the standard integration pipeline outlined by the R package Seurat (v3.2.0, (Butler et al., 2018)). Here, individual temporal samples were normalized independently using default parameters via Seurat::NormalizeData. The log-normalized expression values are then z-scored on the integrated object via Seurat::ScaleData using default parameters after finding anchoring cells between samples.

Dimensional reduction, and cell classification

Choosing an optimal number of principal components (PCs) for dimensional reduction was determined by scree plot using Seurat::ElbowPlot. We selected PCs showing the greatest variance explained until each subsequent PC showed little to no change. For the macrophage time course, we specified 50 total number of PCs to compute and selected the first 49 PCs based on the scree plot to build a shared nearest neighbor (SNN) graph using Seurat::RunPCA and Seurat::FindNeighbors, respectively. Seurat::FindClusters was used with a resolution of 0.8, resulting in 31 clusters. To visualize cells in two-dimensional latent space, we used UMAP dimensional reduction technique via Seurat::RunUMAP using the first 49 PCs aforementioned. For the *il10ra* dataset, we specified 50 total number of PCs to compute and selected the first 28 PCs based on the scree plot to build a shared nearest neighbor (SNN) graph using Seurat::RunPCA and Seurat::FindNeighbors, respectively. Seurat::FindClusters was used with a resolution of 0.8, resulting in 30 clusters. To visualize cells in two-dimensional latent space, we used UMAP dimensional reduction technique via Seurat::RunUMAP using the first 28 PCs aforementioned.

Classification of cell clusters for the macrophage time course and *il10ra* sibling versus mutant dataset were annotated by calculating differential marker expression via Seurat::FindAllMarkers using default parameters.

DE analysis or primary analysis

To distinguish differentially expressed cluster marker genes for both the macrophage time course and the *il10ra* dataset, we used Seurat::FindMarkers to compare cells in one query time point against all other cells (TableS1, TableS3). For all differentially expressed gene tables, we defined our statistical test using Wilcoxon Rank-test. Only genes with a p-value less than 0.05 were retained. Venn diagrams for TableS2 were generated using Manteia (*6*).

HCR-FISH

Hybridization chain reaction (HCR) was performed according to manufacturer's instructions for *il10ra*-B2 (4pmol), *fgl2a*-B2 (2pmol), *irg1*-B2 (2pmol), *mcamb*-B2 (2pmol), *f13a1b*-B2 (2pmol), *tspan10*-B2 (2pmol), *runx3*-B2 (2pmol), *eomesa*-B2 (2pmol), *hepacam2*-B2 (2pmol) and *dusp1*-B2 (2pmol) (*7*) (Molecular Instruments) except that probes were incubated for 48h and amplifiers incubated for 48h. The amplifiers used were B2-546 and B2-647 (Molecular Instruments). HCR fish were subsequently stained with DAPI (5ug/mL) for 30 min at room temperature (RT) in the dark and washed three times with 5x SSCT before imaging. Quantification was made using FIJI (*8*) in the 3D z-Stack.

Time-lapse and confocal imaging

Images were acquired using a Nikon Ti Eclipse with Yokogawa CSU-W1 spinning disk head equipped with a Hamamatsu Flash 4.0 sCMOS. Objective lenses used were a Nikon Plan Apo 40x 1.15NA LWD (water) and a Nikon Plan Apo 20x 0.75NA.
For live imaging experiments, larvae were immobilized with tricaine (MS-222) up to 150 mg/L and mounted in glass bottom dishes (MatTek) with 0.8% low melting point agarose dissolved in 0.5x E2 with tricaine (100 mg/L). Time lapse recordings were started 10-minute after addition of neomycin (300µM) on top of the agarose. Temperature was kept constant at 28.5°C using a Stage Top Chamber (OkoLab).
A Nikon LUNV solid state laser launch was used for lasers 405, 445, 488, 561, and 647nm. Emission filters used on the Nikon were 480/30, 535/30, 605/70.
All image acquisition was performed using Nikon Elements AR 4.6 (Nikon) software.

Macrophage recruitment assay

5dpf larvae expressing *mpeg:GFP* or *irg1*:*lck-mScarletI* or *stat1b*:*lck-mScarletI* and *she*:*lckmScarletI* or *she*:*GAP43-GFP* were treated with 300µM of neomycin for 30min and fixed in 4% PFA 1h, 3h, 5h and 7h after. Larvae were subsequently imaged using spinning disk confocal microscopy. Quantifications of macrophages inside the neuromast was performed in 3D in Imaris 9.0.

Macrophage photoconversion

A circular region of interest was drawn on the neuromast 1h after neomycin to only photoconvert effector macrophages. Photoconversion was performed using the FRAP module of the Nikon spinning disk confocal. We used a 405 laser at 3% and a dwelling time of 300μs. Stimulation was repeated twice to achieve a good photoconversion. The photoconversion is expected to be 30-40% of the protein. The rest will be photoactivated (increased green fluorescence) and will eventually bleach. Then each photoconverted neuromast was imaged with the 445, 488 and 561nm lasers with a z-stack constant of 50μm from the middle of the neuromast and a zStep of 1um (51 slices) with a 40xLWD 3h and 5h after neomycin. Quantification was performed in Imaris 9.0

Macrophage distance and velocity quantifications

Distances and velocities were extracted from time-lapse recordings of *mpeg*:*GFP* and *she*:*lck-mScarletI* 5dpf larvae. The center of the neuromast as well as individual macrophages were tracked over time using Imaris 9.0. Both values were exported form the statistics module of Imaris. Graphs were made using GraphPad Prism 8 (version 8.4.3).

Drug Treatment for GR inhibition

5dpf *mpeg:GFP* larvae were pre-treated with 10μM RU486 (SIGMA M8046) in 0.01%EtOH/0.5xE2 or 0.01%EtOH/0.5xE2 (control) for four hours prior to neomycin treatment. Larvae were kept in RU486 during and after neomycin treatment until fixation in 4%PFA at 4C.

GO and pathway analysis

Gene ontology (GO) and pathway enrichment analysis was performed using Metascape (*9*).

Statistical analysis

All statistical tests were performed using GraphPad Prism 8 (version 8.4.3) as indicated in the figure legends. When comparing data from more than two groups, statistical significance was calculated using one-way ANOVA with Tukey's post hoc test. Data from two groups were compared using two-tailed unpaired t-test. *p-value*s smaller than 0.05 were considered to be statistically significant. Plots were made in GraphPad Prism 8.

Fig. S1

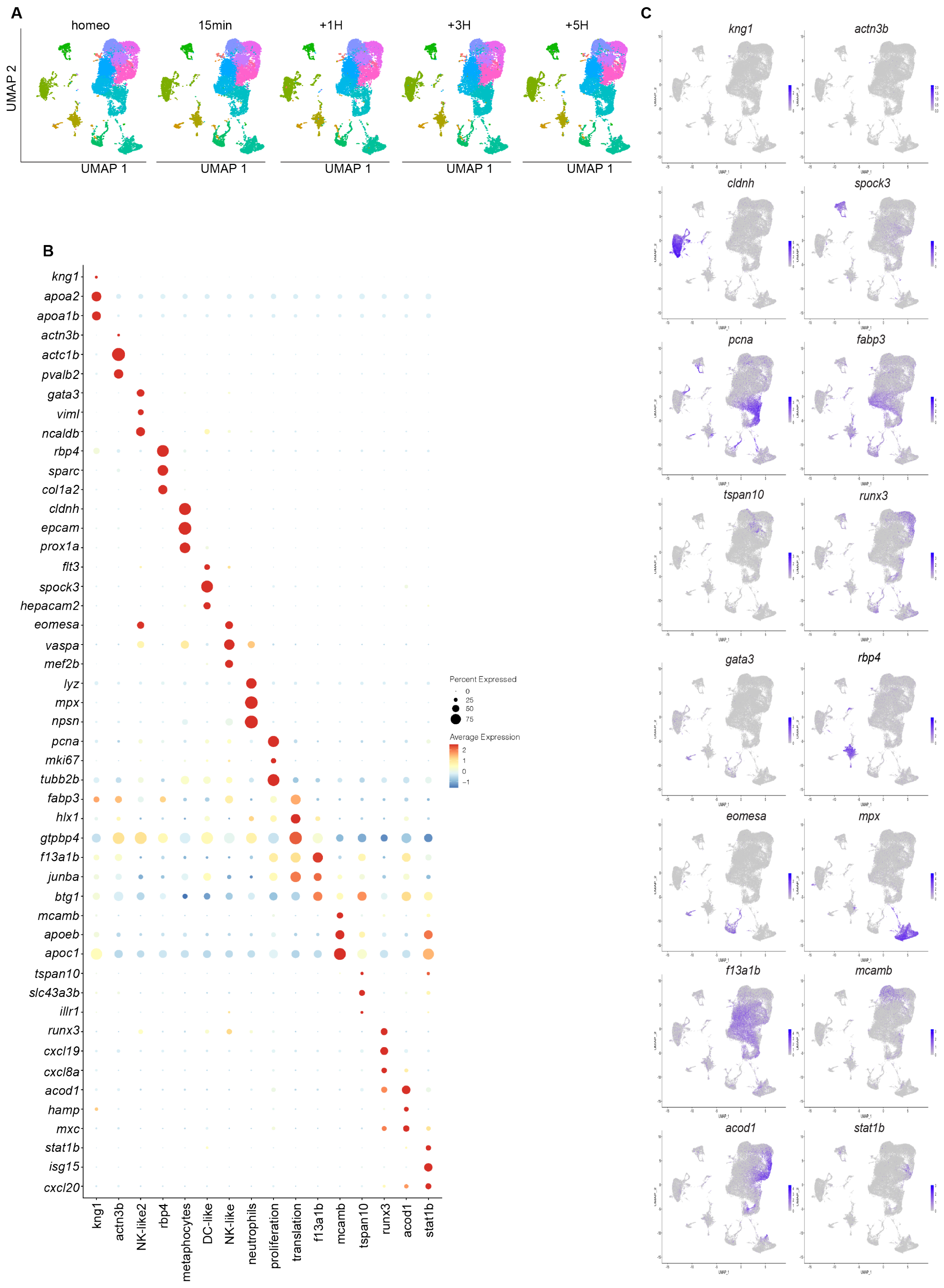
A
homeo
15min
+1H
+3H
+5H
UMAP 2
UMAP 1
B
kng1
apoa2
apoa1b
actn3b
actc1b
pvalb2
gata3
viml
ncaldb
rbp4
sparc
col1a2
cldnh
epcam
prox1a
flt3
spock3
hepacam2
eomesa
vaspa
mef2b
lyz
mpx
npsn
pcna
mki67
tubb2b
fabp3
hlx1
gtpbp4
f13a1b
junba
btg1
mcamb
apoeb
apoc1
tspan10
slc43a3b
illr1
runx3
cxcl19
cxcl8a
acod1
hamp
mxc
stat1b
isg15
cxcl20
kng1
actn3b
NK-like2
rbp4
metaphocytes
DC-like
NK-like
neutrophils
proliferation
translation
f13a1b
mcamb
tspan10
runx3
acod1
stat1b
Percent Expressed
0
25
50
75
Average Expression
2
1
0
−1
C
kng1
actn3b
cldnh
spock3
pcna
fabp3
tspan10
runx3
gata3
rbp4
eomesa
mpx
f13a1b
mcamb
acod1
stat1b

**Fig. S1. Cluster markers from the macrophage scRNA-seq time-course. (A)** Individual UMAP for each dataset. 14000 cells per dataset. (**B**) DotPlot showing three marker genes per cluster. (**C**) Feature plots of cluster marker genes.

**Fig. S2**

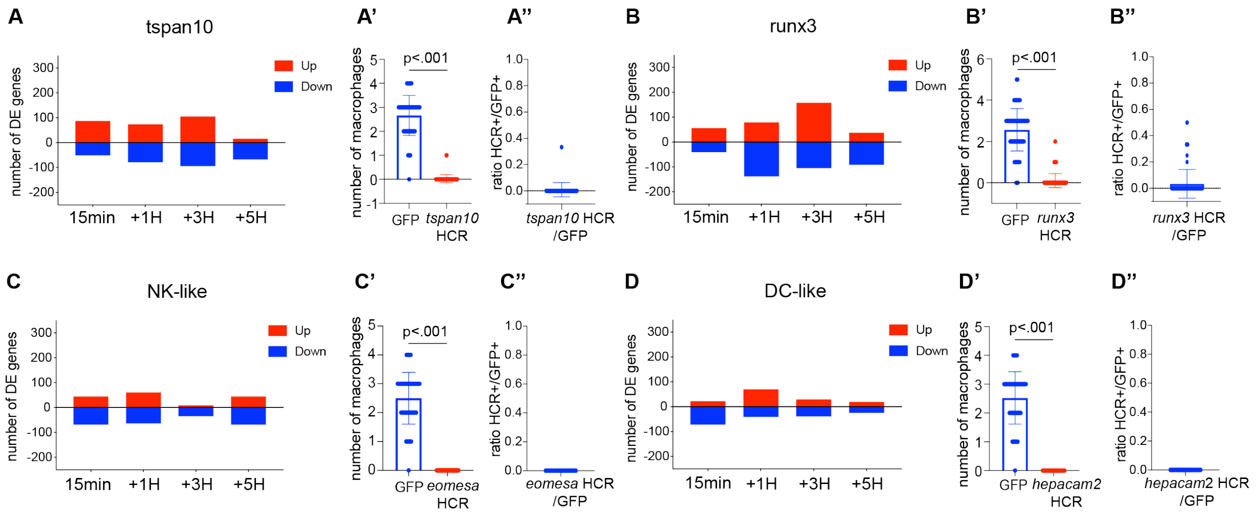


**Fig. S2. Even non-effector macrophages show gene expression changes. (A, B, C and D)** Quantification of genes differentially up- and downregulated at each timepoints. (**A', B', C' and D'**) Quantifications of GFP+ effector cells and effector cells with a positive HCR signal. Each dot represents the number of macrophages per neuromast (5 neuromasts per larvae and 16 larvae). P-values represent non-parametric Student t-test. (**B''and D''**) Quantifications of the ratio between HCR+ cells and GFP+ effector macrophages.

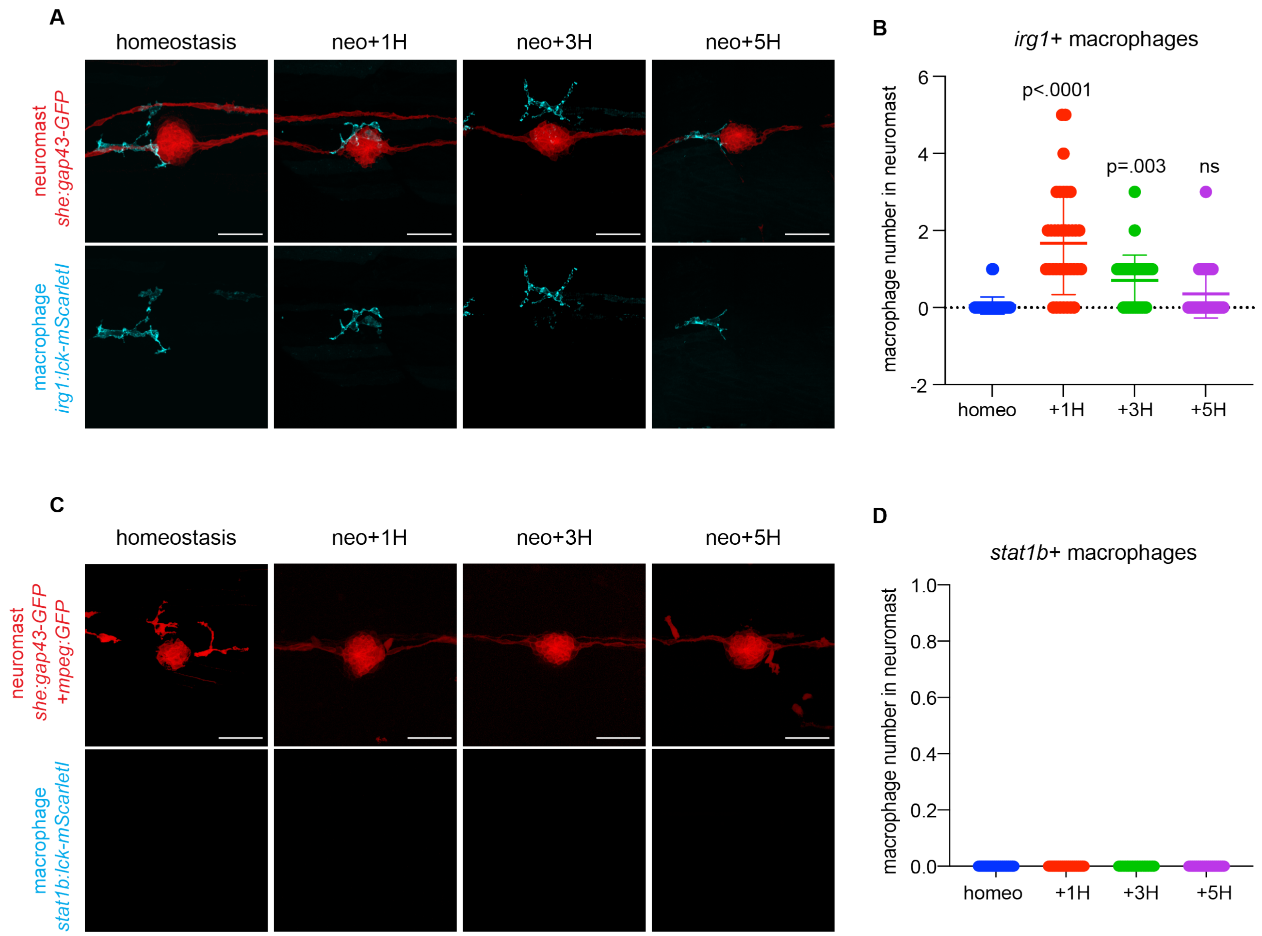


**Fig. S3. 'irg1/acod1' but not 'stat1b' macrophages are effector cells. (A, C)** Representative confocal images (projection of a 30µm z-stack) of the macrophage recruitment assay. **(B, D)** Quantification of macrophages inside the neuromasts. Each dot represents the number of macrophages per neuromast (3 neuromasts per larvae and 6 larvae per condition). P-values represent a post-hoc (Tuckey) test from each condition relative to homeostasis.

**Fig. S4**

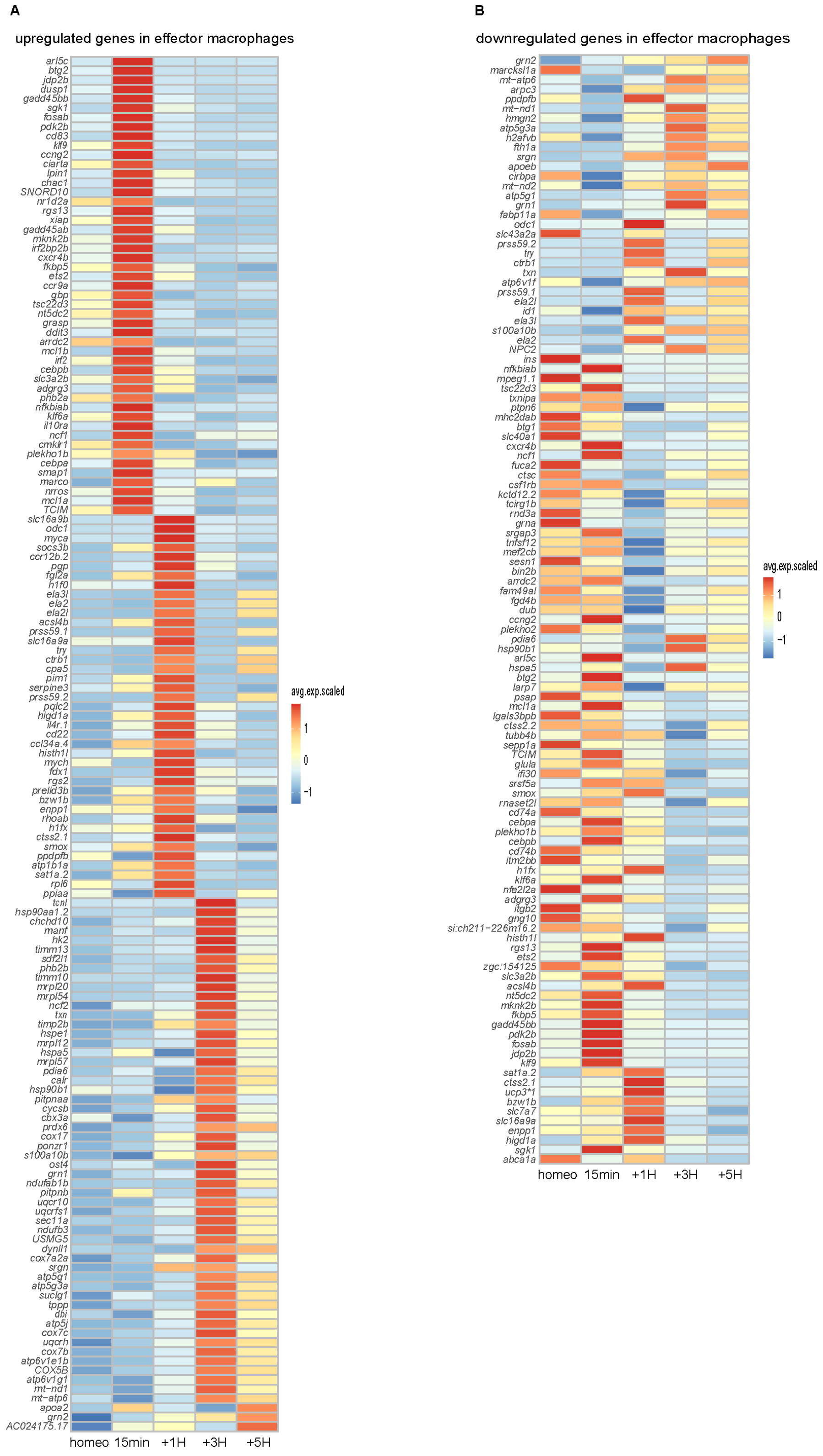


**Fig. S4. Differentially expressed genes at each timepoint. (A-B)** Heatmaps for (A) up- and (B) downregulated genes at each timepoint.

Fig. S5

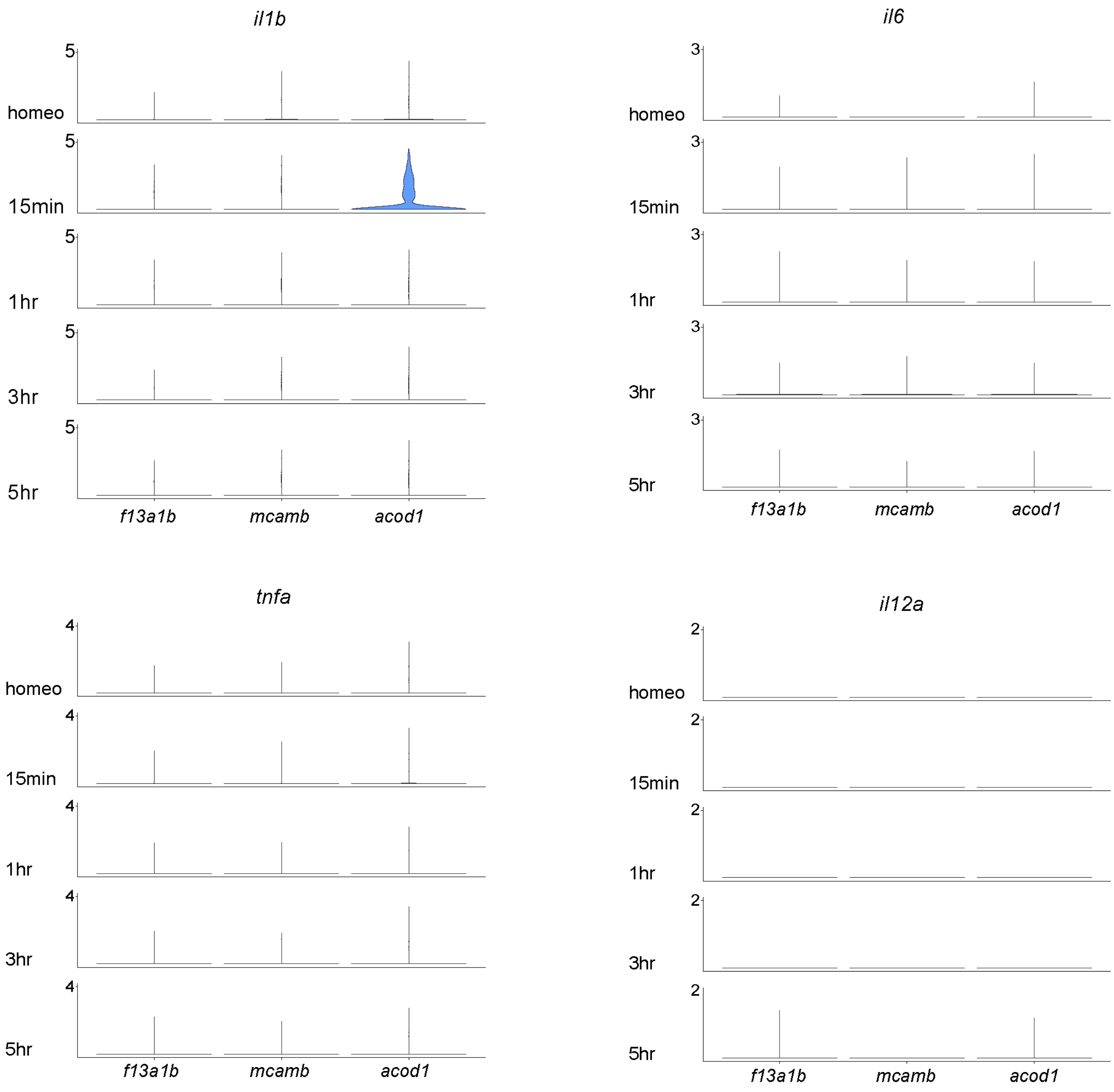


**Fig. S5 Most pro-inflammatory cytokines are not transcriptionally upregulated in effector macrophages after HC death.** Stacked Violin-Plots for *il1b*, *il6*, *tnfa* and *il12a.*

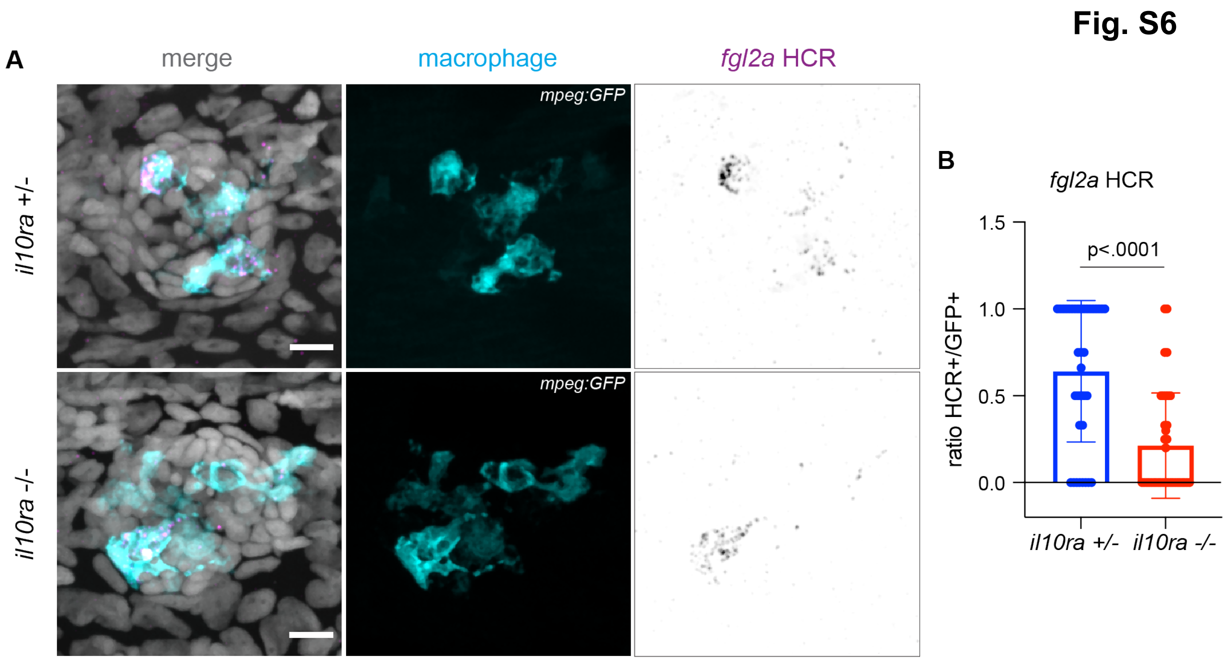


**Fig. S6 The IL10 signaling target gene *fgl2a* is downregulated in the *il10ra* mutant.** (**A**) Representative confocal images (projection of a 30µm z-stack) of HCR-FISH within the effector macrophages for f*gl2a.* (**B**) Quantifications of the ratio of HCR+ cells over GFP+ effector macrophages for f*gl2a* in *il10ra* mutant heterozygous and homozygous (n=36 neuromasts from 12 larvae per condition). P-values represent non-parametric Student t-test.

**Fig. S7**

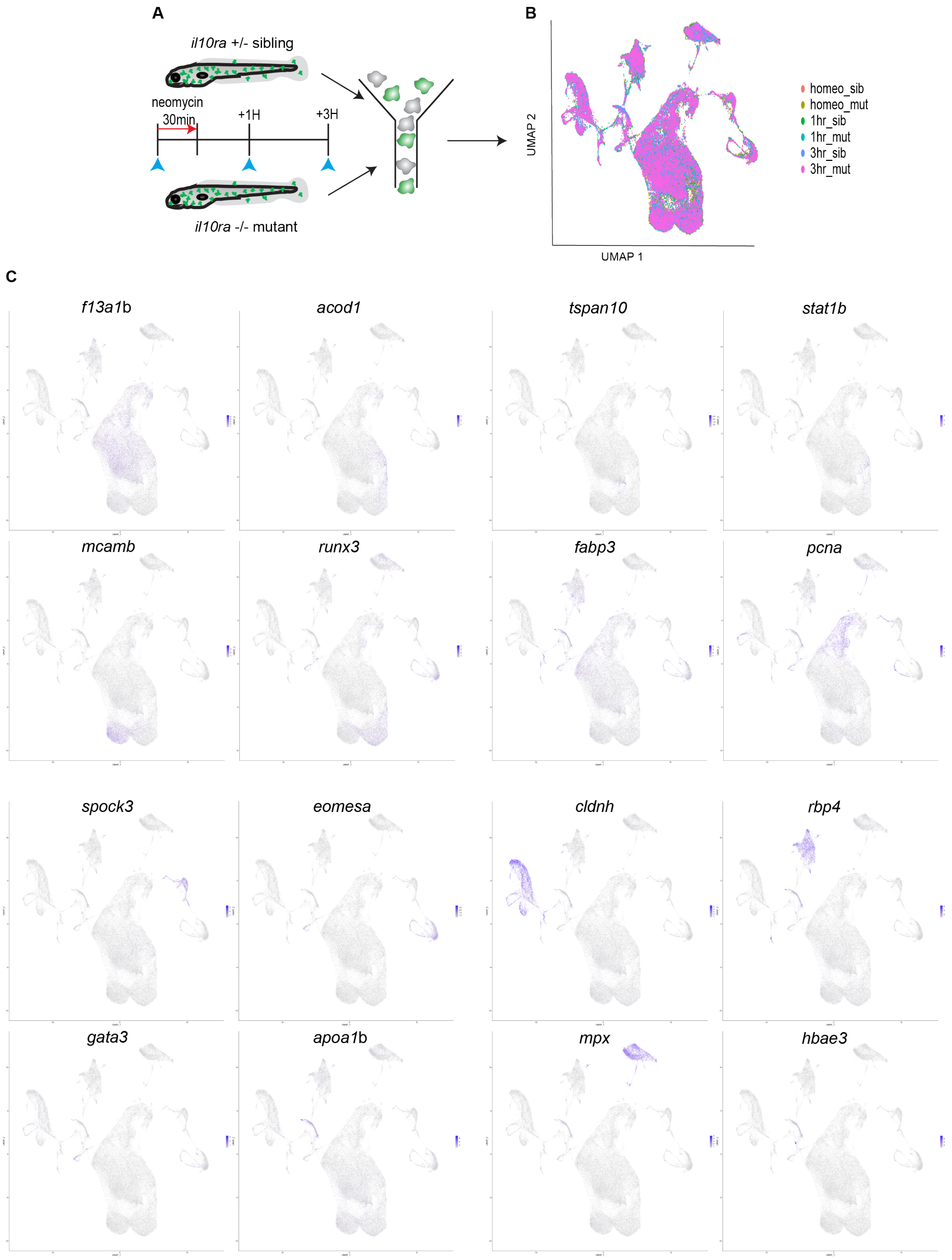
A
il10ra +/- sibling
neomycin
30min
+1H
+3H
il10ra -/- mutant
B
UMAP 2
UMAP 1
homeo_sib
homeo_mut
1hr_sib
1hr_mut
3hr_sib
3hr_mut
C
f13a1b
acod1
tspan10
stat1b
mcamb
runx3
fabp3
pcna
spock3
eomesa
cldnh
rbp4
gata3
apoa1b
mpx
hbae3

**Fig. S7 Cluster markers for the *il10ra* mutant macrophage in the scRNA-seq time-course.** **(A)** Schematics of neomycin regime and timepoint collection for scRNA-seq. (**B**) Integrated UMAP of the 6 datasets. (**C**) Feature plots of cluster marker genes.

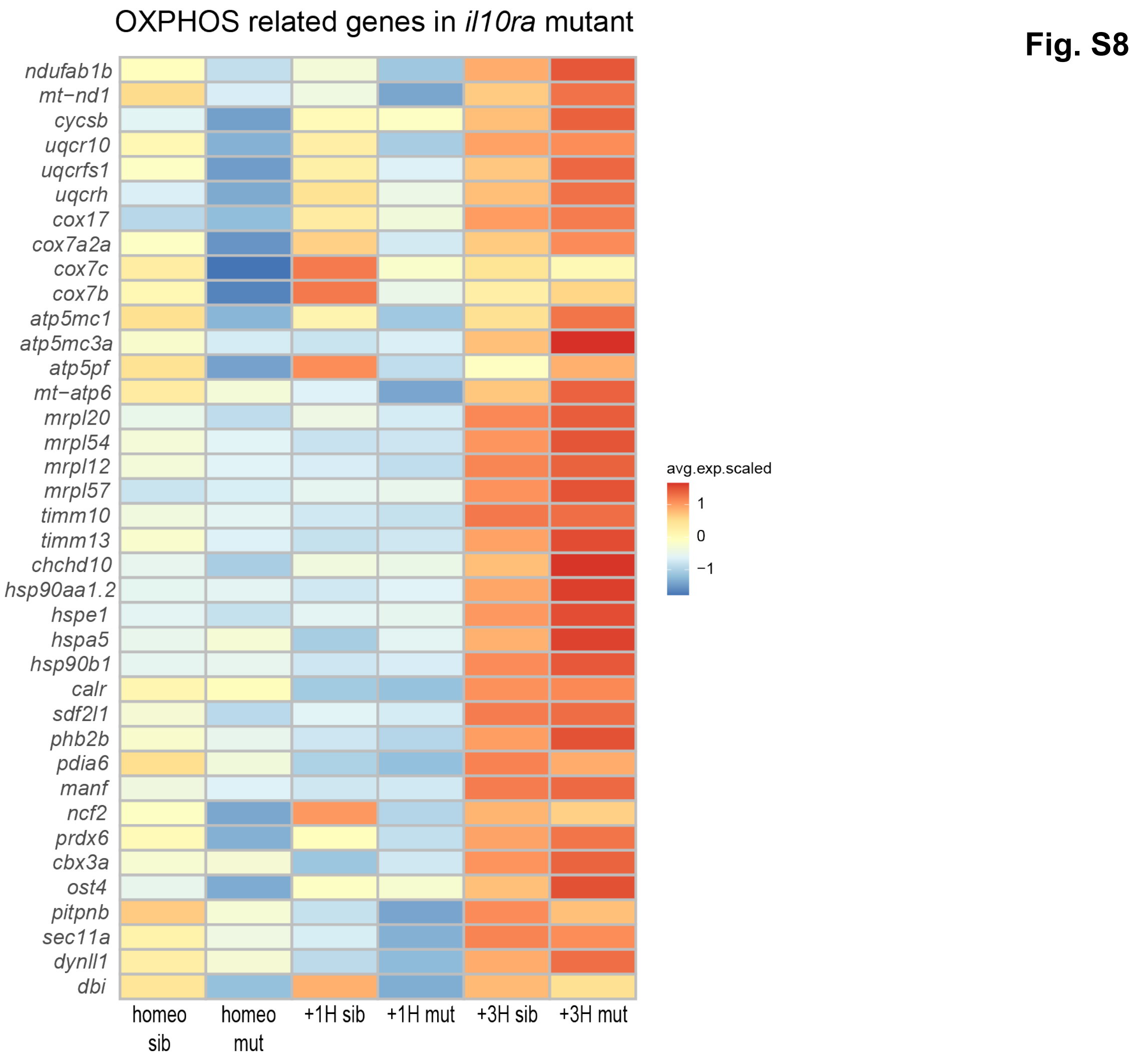


**Fig. S8 Induction of oxidative phosphorylation related genes is not affected in the *il10ra* mutant.** Heatmap for oxidative phosphorylation related genes at each timepoint between *il10ra* mutant and siblings from the scRNA-seq datasets.

**MovieS1 3D animation of a whole 5dpf larva related to Figure1A.** Macrophages (*mpeg:GFP*) are labelled in cyan and neuromasts (*she:lckmScarletI*) in red.

**MovieS2 Effector macrophages rapidly phagocytose dying HCs upon addition of neomycin.** Maximum projection (z-stack of 30µm) of a time-lapse recording of macrophages (blue, *mpeg*:*GFP*) and HCs (red, *myo6*:lckmScarletI). One image is taken every minute for two hours.

**MovieS3 Zoom of macrophages phagocytosing HCs from MovieS2**

**MovieS4 Dynamics of macrophages during 7H after neomycin treatment related to Figure1B.** Maximum projection (z-stack of 30µm) of a time-lapse recording of macrophages (cyan, *mpeg*:*GFP*) and neuromasts (red, *she*:*lckmScarletI).* One image is taken every five minutes for seven hours.

**MovieS5 Effector macrophages are located close to the neuromasts, and all macrophages increase their velocity upon neomycin treatment.** Maximum projection (z-stack of 90µm) of a time-lapse recording of macrophages (cyan, *mpeg*:*GFP*) and neuromasts (red, *she*:*lckmScarletI*) in the trunk region of 5dpf larvae. One image is taken every minute for two hours.
